# MicroED Structures of Fluticasone Furoate and Fluticasone Propionate Provide New Insights to Their Function

**DOI:** 10.1101/2024.09.18.613782

**Authors:** Jieye Lin, Johan Unge, Tamir Gonen

## Abstract

The detailed understanding of fluticasone, a widely prescribed medicine for allergic rhinitis, asthma, and chronic obstructive pulmonary disease (COPD), has not been complete due to challenges in structural elucidation. The three-dimensional (3D) structure of fluticasone furoate **1** remained undetermined for decades, while the existing structures of fluticasone propionate **2** required refinement against improved data. In this study, we applied microcrystal electron diffraction (MicroED) to determine the 3D structures of **1** and **2** in their drug formulation state. Density functional theory (DFT) calculations were utilized to model solvent effects to determine the preferred geometries in solution. A comparative analysis of structures of **1** and **2** across three states (drug formulation state, in solution, and biologically active state) revealed major conformational changes during the entire transition. Potential energy plots were calculated for the most dynamic bonds, uncovering their rotational barriers. This study underscores the combined use of MicroED and DFT calculations to provide a comprehensive understanding of conformational and energy changes during drug functioning in humans. The quantitative comparison highlights the subtle structural differences that can lead to significant functional changes in pharmaceutical properties.

## 1 Introduction

Fluticasone is widely used to treat allergic rhinitis, asthma, and chronic obstructive pulmonary disease (COPD).^1-3^ In 2021, it was ranked as 23^rd^ most-prescribed medicine in the United States, with nearly 8 million patients and 25 million prescriptions per year.^4^ The common forms of fluticasone are fluticasone furoate **1** and fluticasone propionate **2**. Depending on the pharmaceutical formulation, they are marketed under different brand names, such as Flonase Sensimist,^5^ Arnuity Ellipta,^6^ Flovent Disku,^7^ Cutivate.^8^

Characterizing the three-dimensional (3D) structures of **1** and **2** in their drug formulation state has been of a long-term interest. Previous research has shown that **1** co-crystalized with multiple guest molecules (Table S1, Supporting Information).^9-12^ Five of these co-crystals formed large single crystals, and were solved by single-crystal X-ray diffraction (SC-XRD), however the structures were not disclosed to the scientific community; most solvomorphs of **1** were tiny microcrystals that did not allow structure determination using SC-XRD and could only be indexed by powder X-ray diffraction (PXRD);^9-12^ unsolvated crystal form of **1** were used in pharmaceutical formulation, however for the known three forms were missing their 3D structures.^9-12^ As for **2**, two polymorph structures have been determined by SC-XRD and PXRD (Table S1, Supporting Information).^13-15^ The form1 SC-XRD structure (CSD entry: DAXYUX)^14^ showed disorder in the 17β-fluoromethylthioester moiety and a relatively high R_1_ value (7.5 %); the form2 was specifically obtained from supercritical crystallization, and its PXRD structure (CSD entry: DAXYUX01) was missing hydrogen atoms.^15^ The unsolvated crystal structure of **1** remained elusive up to this day; the current two polymorph structures of **2**, especially form1 seemed to benefit from improved data. The development of microcrystal electron diffraction (MicroED) techniques bypassed the crystal size limitation for SC-XRD and is particularly suitable for micro- or nano-sized crystals, *i*.*e*. crystals are only a billionth of the size commonly used in SC-XRD.^16,17^ In this study, we applied MicroED to unveil the 3D structures of **1** and **2**. Not merely a supplement to the literature, these long awaited crystal structures reveal the conformations of **1** and **2** in the drug formulation state, representing the “starting point” towards their active conformation in humans.

Chemically, both **1** and **2** exhibit a consistent steroidal backbone with identical substitution groups; the only difference is the 17α-esterification, with a furoate ester in **1** and a propionate ester in **2**. This similarity in chemistry allows **1** and **2** to perform the same biological function, targeting the glucocorticoid receptor (GR) as agonists.^18,19^ Upon binding, the GR complex undergoes conformational changes and translocation into nucleus, which can modulate gene expression.^20,21^ The binding affinity to GR differs between **1** and **2**, with **1** being nearly 70% stronger than the **2**.^1,22^ Clinically, **1** demonstrates a faster association rate and a slower dissociation rate compared to **2**; the daily dose requirement for **1** is 110 μg, smaller than the 200 μg for **2**.^1^ The complex structures for GR/**1** and GR/**2** have been solved by X-ray crystallography and CryoEM (GR/**1**, PDB entries: 3CLD, 7PRV; GR/**2** was not disclosed).^18,19^ Structural analysis showed comparable residue interactions of GR/**1** and GR/**2**, but a better fit of 17α-pocket for furoate ester in **1** than propionate ester **2**.^18^ The rationalizing of the association and dissociation rate differences for **1** and **2** is not adequate without understanding their conformational and energy changes upon binding to the target protein (biologically active state). A smaller conformational change may be related to a smaller energy barrier and lead to a faster association. A larger flexibility on the other hand is related to an increase in entropy upon dissociation and a more favorable release. It may be speculated whether a lower energy barrier also facilitates the release of the compound from its binding position if there is a large difference between the bound and free form of the compound.

The solution structure serving as a “transition form” for pharmaceuticals is presumably more related to the biologically active conformation and therefore undergoes a lower energy barrier to its final form; however, such conformations are numerous in equilibrium and difficult to model. In this study, we applied density functional theory (DFT) calculations to model solvent effects and the preferred geometries of **1** and **2** in water. Then, three states of **1** and **2** (drug formulation state, in solution, and biologically active state) were compared to show the major conformational changes. Potential energy plots for the most dynamic bonds were calculated, uncovering their rotational barriers during the transition from drug formulation state to biologically active state, allowing to quantitatively explain the structure-function differences between **1** and **2**.

## 2 Results and Discussion

The commercially purchased **1** and **2** were recrystallized from methanol at room temperature, forming needle-shaped microcrystals on the surface of glass vials (Figure 1). Recrystallization was assumed to result in the unsolvated **1** and **2** following the procedure described in the literature.^11,12,23^ The crystals were gently ground into fine powders using a spatula. The MicroED grid preparation followed the procedure described in the literature (See “Methods” in Supporting Information).^24^ The TEM grids containing microcrystals of **1** and **2** were loaded in a 200 keV Talos Arctica Cryo-TEM (Thermo Fisher). Crystals that were of suitable thicknesses (with a preferred image contrasts) were selected under the imaging mode (SA 3400×) and calibrated to their eucentric heights to maintain them within the beam area during the continuous rotation.

**Figure 1.**
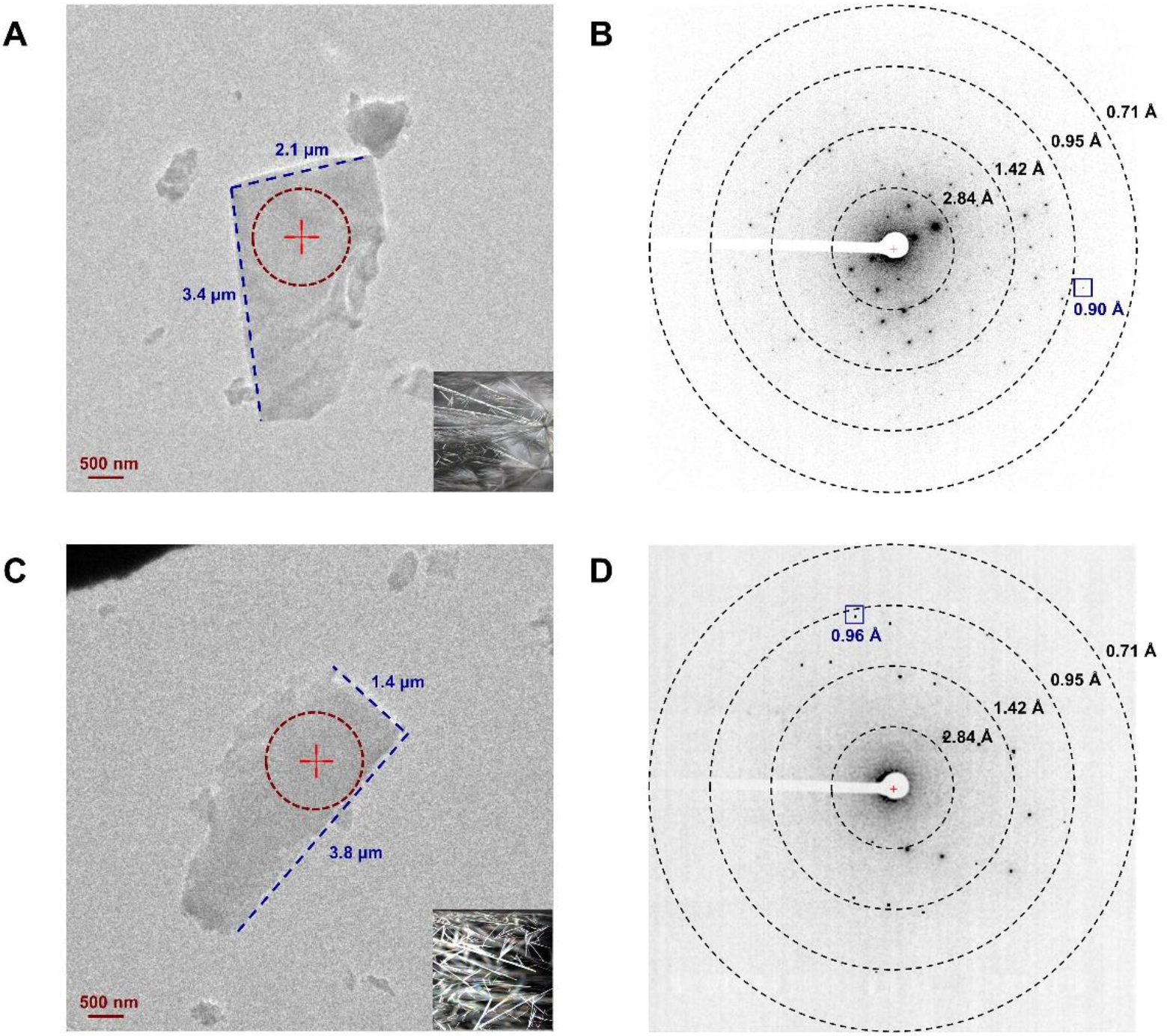
View of the crystals and their diffraction patterns. (A, C) Images of **1** and **2** under imaging mode (SA 5300×) and the stereo microscope (16×), respectively. (B, D) Diffraction patterns of **1** and **2** under diffraction mode (659 mm), respectively. The diffraction beam area was highlighted in dashed red circles in Figures 1A and 1C.

MicroED data were collected under the diffraction mode with a camera length of 659 mm (the calibrated sample-detector distance). Typical data collection used a 0.5 s exposure time and 2° per second rotation rate settings over 120° wedges (−60° to +60°), which can be collected in ∼1 min with a total dose of ∼0.60 e^-1^/Å^2^ (electron dose rate: ∼0.01 e^-1^/(Å^2^·s)).^25^ The extended rotation wedge captured high-tilt diffraction data to increase the completeness. The starting and ending angles were carefully truncated to avoid overlapping with nearby crystals or the grid bar.

MicroED data were saved in mrc format and converted to smv format using mrc2smv software (https://cryoem.ucla.edu/microed).^25^ The converted frames were indexed, integrated, and scaled in XDS.^26,27^ **1** was indexed with orthorhombic space group P 2_1_2_1_2_1_ (**a**=7.70 Å, **b**=13.95 Å, **c**=23.48 Å, **α**=90.0°, **β**=90.0°, **γ**=90.0°), and one dataset only was enough to reach ∼96% in completeness; **2** was indexed with monoclinic space group P 2_1_(**a**=7.57 Å, **b**=14.06 Å, **c**=10.86 Å, **α**=90.0°, **β**=99.4°, **γ**=90.0°), with four datasets merged to achieve ∼95% completeness (Table S2, Supporting Information). The intensities were converted to SHELX hkl format using XDSCONV,^27^ and were directly solved by SHELXD^28^ at the resolution of 0.90 Å and 0.96 Å for **1** and **2**, respectively. The structures were then refined by SHELXL,^29^ reaching the lowest R_1_ values of 16.4% and 15.3% for **1** and **2**, respectively (Table S2, Supporting Information). The non-hydrogen atoms were accurately determined from the potential maps at sub-atomic resolution for **1** and **2** (Figures 2C-D). Both structures were unsolvated and devoid of methanol molecules. The 17β-fluoromethylthioester moiety in **1** showed no signs of disorder (Figure 2C), and the furoate ring oxygen atom (O6) was carefully examined by comparing the measurements of adjacent C‒O (1.41 Å) and C=C (1.32 Å) bond lengths to their reference bond lengths.^30^ MicroED structure of **2** (Figure 2D) matched with the previously determined X-ray structure of **2** (CSD entry: DAXYUX), which contained a disordered fluoromethyl group^14^ The polar H atoms were located in the omit map, while the non-polar H atoms were placed using riding models.

**Figure 2.**
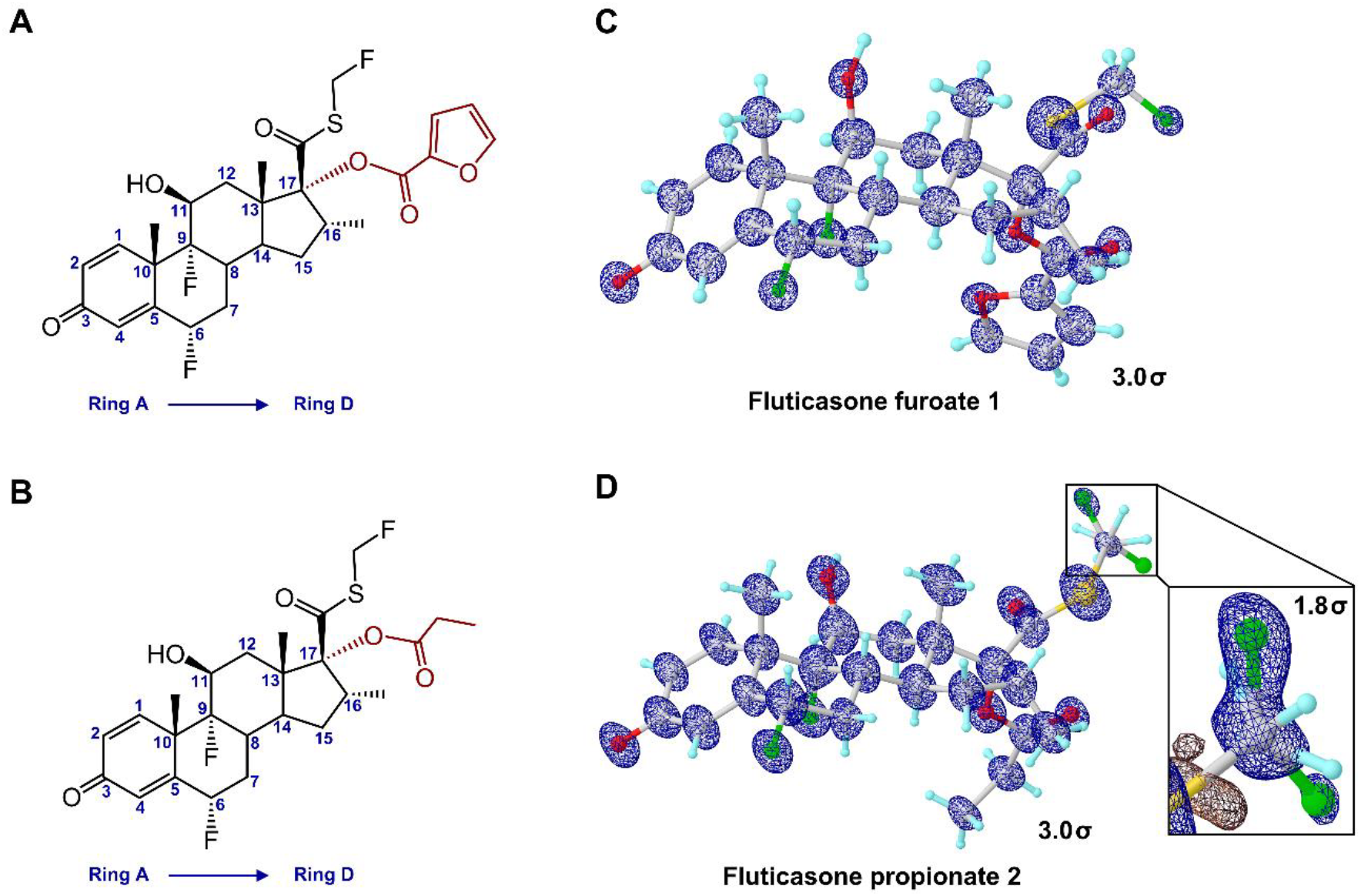
Chemical and MicroED structures. (A, B) Chemical structures of **1** and **2**, respectively. (C, D) MicroED structures of **1** and **2**, respectively. 2Fo-Fc density maps (3-sigma) were shown in blue mesh. The density of minor conformation of fluoromethyl group in **2** was shown in expanded box at 1.8-sigma.

Atoms were numbered following the steroid numbering convention described in literature (Figures 2A-B; Figure S1, Supporting Information).^31^

Crystal packing was compared for MicroED structures **1** and **2**. In **1**, molecules were tightly packed via a repetitive hydrogen bond O2‒H^**…**^O1 (2.71 Å) between 11β-hydroxy group (O2) and 3-keto group (O1) along the *b*-axis (Figure 3A). A similar hydrogen bond O2‒H^**…**^O1 (2.76 Å) was also found in **2** (Figure 3B). Weak contacts, such as C‒H^**…**^F contacts (H^**…**^F<3.0 Å),^32^ extend the packing along other directions but vary between **1** and **2**. For example, in **1**, C1‒ H^**…**^F2 (2.56 Å) and C25‒H^**…**^F2 (2.93 Å) around 6α-fluorine (F2) can extend crystal packing along *a*-and *c*-axes; while in **2**, three fluorine atoms (F1, F2, and F3A) form at least seven contacts that extend crystal packing along three axes, such as C19‒H^**…**^F3A (3.39 Å), C24‒ H^**…**^F2 (3.41 Å) and C24‒H^**…**^F3A (3.39 Å). The existence of minor conformation of F3B bridges more contacts to C7 and C14. A non-uniform crystal growth with an increased growth along the *b*-axis over the other two directions led to the plate- or needle-shaped morphologies in **1** and **2**. Voids (empty space) in the unit cells of **1** and **2** were further examined and it was found that 8.7% of the unit cell volume (55 Å^3^ per molecule) in **1** is accessible to solvent, whereas 0.8% of unit cell volume (4.5 Å^3^ per molecule) is accessible to water in **2**, indicating a better permeability of water in **1** than in **2**.

**Figure 3.**
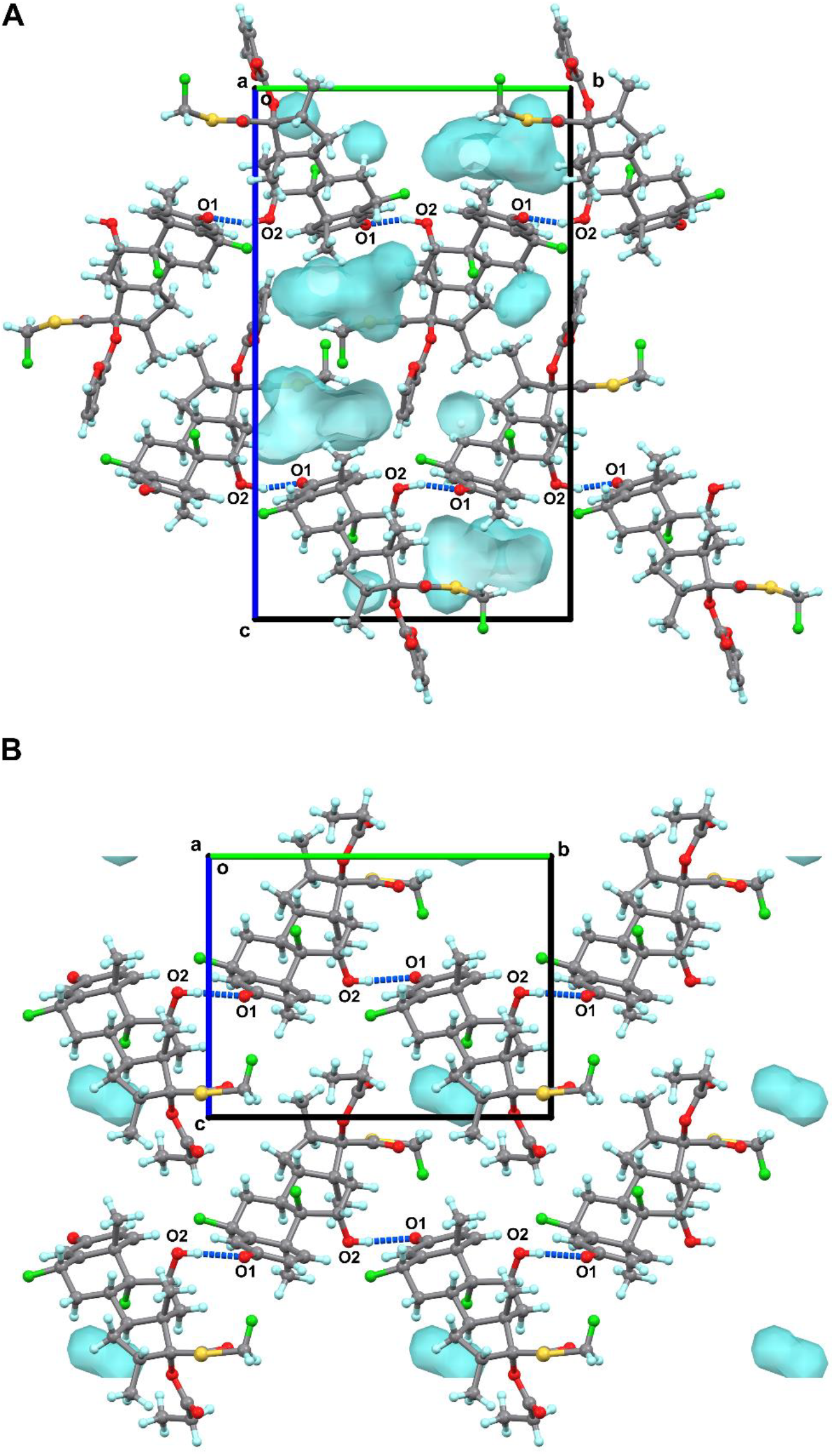
Packing diagram of **1** (A) and **2** (B), viewed along *a*-axis. Hydrogen bonding interactions were represented by the dashed lines in blue. Voids were detected by the same probe radius and grid spacing settings, colored in cyan. The minor conformation of **2** was omitted for clarification.

Examining the structural parameters for the asymmetric units of **1** and **2**, we found that both contain a rigid steroidal backbone composed of four fused rings (rings A-D, from left to right, see Figures 2A-B). These four rings act as conformational constraints for the overall backbone. Only a few neglectable differences were found in the steroidal backbone conformations between **1** and **2**. These minor variations occurred in the fusion bonds of rings B and C but did not affect rings A and D (Figure S1, Supporting Information). The major structural differences between **1** and **2** were observed in 17β-fluoromethylthioester substitution. For example, the C13‒C17‒ C20‒S1 is -74.2° in **1**, while it is 108.5° in **2** (Figure S1, Supporting Information), with nearly a 180° difference. Rotation of C17‒C20 bond leads to more structural hinderances in the extended portion, with substantial rotational barriers. The 17α-esters in **1** and **2** share similar conformations but differ at the terminal side, for instance, the lipophilic part of furoate ring (C25) positions outward due to the interaction with 6α-fluorine (F2) in **1**; whereas the flexible ethyl group (C24) positions backward to contact with both 6α-fluorine (F2) and 17β-fluoromethylthioester moiety (F3A) in **2** (Figure S1, Supporting Information).

The different conformations of the two substitutions, together with their varying rotational barriers required for **1** and **2**, likely results in different free energy changes for **1** and **2** in their transition to their biologically active state from their drug-formulation state. These energy differences are also a result of the interactions within the protein pockets, depending on the exact chemical contexts. This in term influences the rate of transition from the drug formulation state to the biologically active state and significantly influences the pharmaceutical properties, like association/dissociation rate,^33^ and biological half-life.^34^

Solid-state drugs generally need to dissolve in human plasma before interaction with target protein, however the structure in solvent is challenging to model since there is an ensemble of conformations as an equilibrium. Here, we applied density functional theory (DFT) calculation to model solvent effects and the preferred geometries of **1** and **2** in water (See “Methods” in Supporting Information). Geometric optimization was performed using functional/basis set combination B3LYP/6-31G(d,p),^35,36^ with the solvent effects of water modeled by conductor-like polarizable continuum model (CPCM)^37^ and the solvation model based on density (SMD),^38^ both implemented in ORCA 5.0 software.^39^ The B3LYP/6-31G(d,p)^35,36^ optimized structures were further validated by comparing with models calculated from ωB97X/6-311G(d,p),^40,41^ B3LYP/6-311G(d,p)^36,41^ and showed no discernible variances caused by the different functional/basis sets. Comparing the structures of **1** and **2** in their drug formulation state with DFT-calculated structures in solution showed very minor conformational changes in substitution groups (Figure 4). For example, in **1**, the O4‒C22‒C23‒O6 has 16° rotation, twisting the furoate ring from 15.4° to -0.6°; in **2**, the 3-keto group (O2) and propionate ester (C23, C24) exhibit at most a 0.4 Å movement due to molecular stretching. These minor conformational changes suggest both **1** and **2** are highly conformational constrained in their drug formulation state and in solution, allowing them to maintain a consistent geometry during the transition to their biologically active state in the protein pocket.

**Figure 4.**
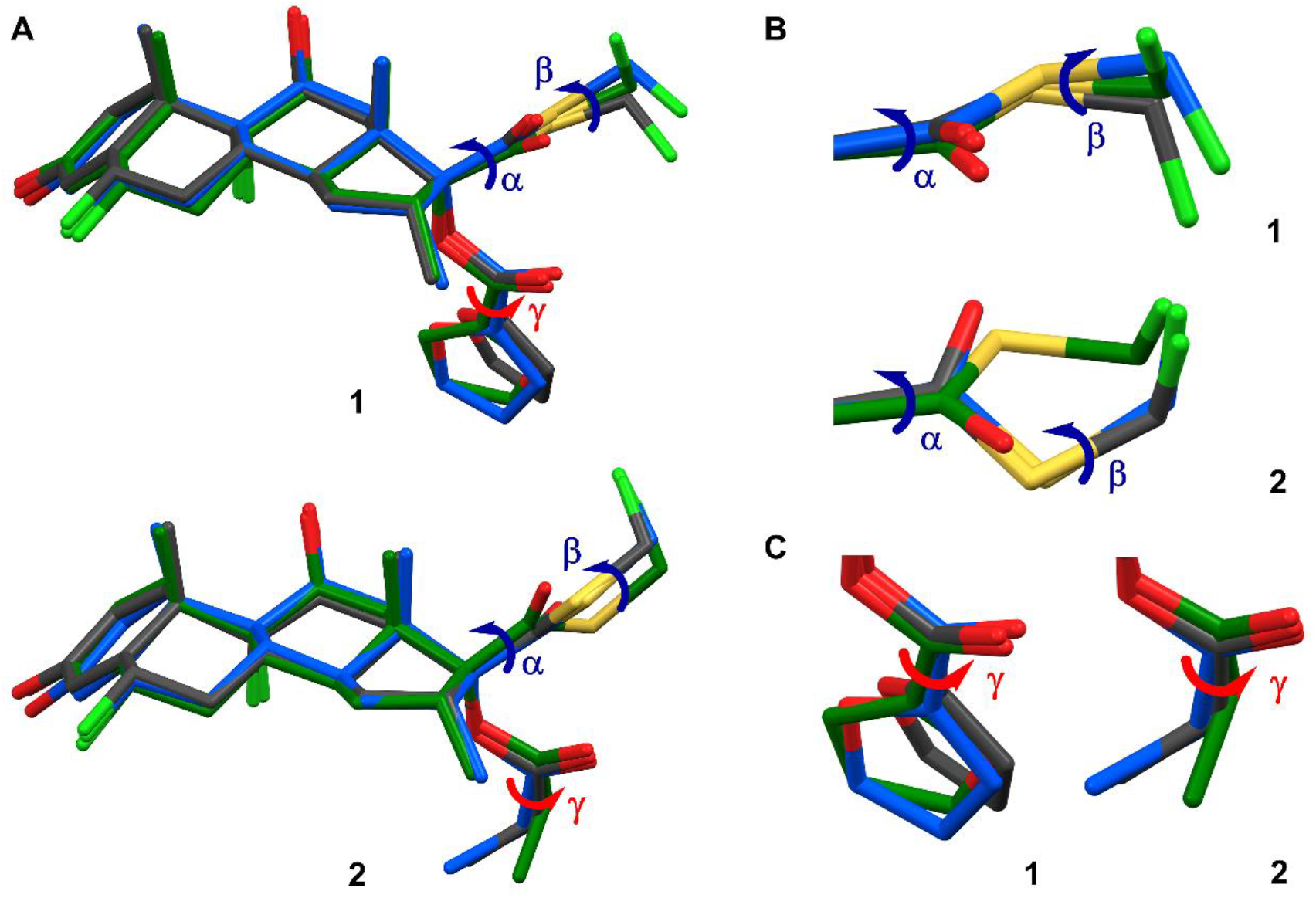
Structure overlays of **1** and **2**. (A) Overlay of structures of **1** and **2** in drug formulation state (grey), in solution (blue), and in biologically active state (green). Three torsion angles **α** (C13‒C17‒C20‒S1), **β** (C20‒S1‒C21‒F3) and **γ** (O4‒C22‒C23‒O6 in **1**, O4‒C22‒C23‒C24 in **2**) were marked representing the major conformational changes from drug-formulation state to biologically active state. A structure model was manually made for **2** in biologically active state according to the presenting in literature.^18^ The minor conformation of **2** was omitted for clarification. (B) Expanded view and comparison of **α** and **β** torsion angles in **1** and **2**. (C) Expanded view and comparison of **γ** torsion angle in **1** and **2**.

Both **1** and **2** act as GR agonists in humans, and their complex structures have been reported (GR/**1**, PDB entries: 3CLD, 7PRV; GR/**2** was presented in literature but was not disclosed).^18,19^ A surrogate structure was manually created herein based on a figure from the literature to mimic the biologically active state of **2** (Figure 4).^18^ To understand the conformational changes when **1** and **2** transition from drug formulation state to biologically active state, the structures of **1** and **2** in three states (drug formulation state, in solution, and biologically active state) were compared (Figure 4). Three torsion angles **α** (C13‒C17‒C20‒S1), **β** (C20‒S1‒C21‒F3) and **γ** (O4‒C22‒ C23‒O6 in **1**, O4‒C22‒C23‒C24 in **2**) were identified as responsible for the major conformational changes. Notably, the **β** and **γ** torsion angles in **1**, and the **α** and **β** torsion angles in **2**, exhibited nearly 180° changes from drug formulation state to biologically active state (Figures 4B-C). To quantitatively model the rotational barrier and energy landscapes, the relative potential energy plots were calculated by scanning **α, β** and **γ** torsions every 15° from 0° to 360° (See “Methods” in Supporting Information). The values of **α, β, γ** torsion angles and the corresponding potential energies in **1** and **2** were highlighted and compared (see points “a-c” in Figure 5).

**Figure 5.**
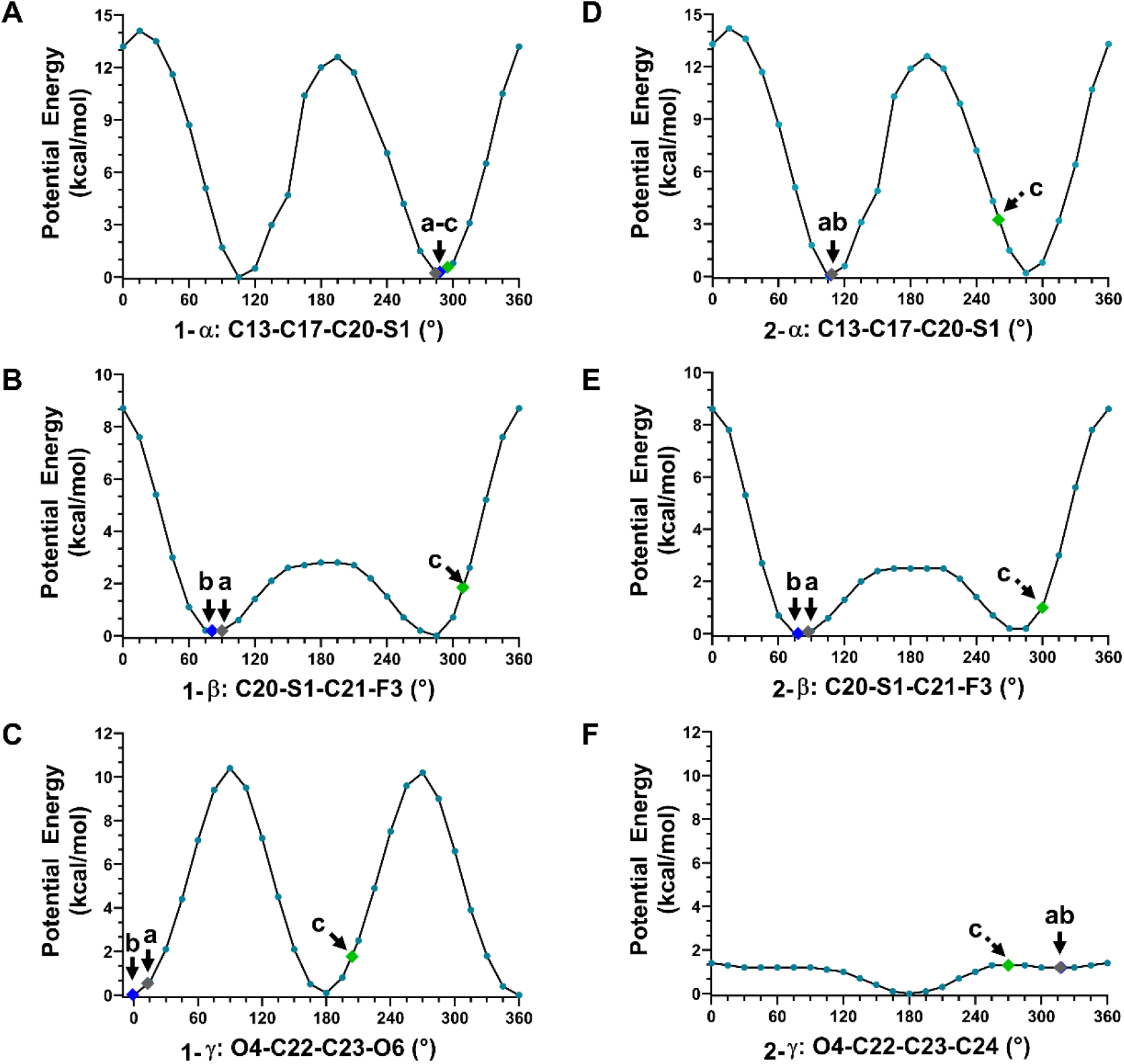
DFT-calculated potential energy plots for **1** (A-C) and **2** (D-F), showing the potential energy changes caused by a rotation of the corresponding torsions. Three points “a-c” were highlighted to represent the corresponding torsion angles measured from drug formulation state (point a), in solution (point b), biologically active state (point c).

The rotation of **α** torsion angle significantly affects the overall conformation of 17β-fluoromethylthioester moiety and involves a rotational barrier of ∼14 kcal/mol (Figures 5A and 5D). In the biologically active state, the O3 atom of the carbonyl group is positioned outward, either hydrogen bonded to Cys736 residue (Figure S3B, Supporting Information)^19^ or hydrophobically interacted with Tyr735 residue (Figure S3A, Supporting Information).^18^ Due to the large rotational barrier, it is energetically unfavorable to rotate the **α** torsion angle. In **1**, the **α?**is relatively fixed, with 284° in drug formulation state, 288° in solution, and 295° in biologically active state,^18^ resulting in an energy change of less than 1 kcal/mol (Figure 5A). On the contrary, **2** undergoes more than 150° rotation in **α**, with 109° in drug formulation state, 108° in solution, and ∼260° in biologically active state.^18^ Although **2** ends with a low energy value (∼3.3 kcal/mol), the transition still requires ∼13 kcal/mol to overcome the rotational barrier (Figure 5D). This may indicate a slower association rate with the receptor for **2** compared to **1**. Although the pathway is complex, it is more likely that unbound 17β-fluoromethylthioester moiety in **2** be free to be metabolized to 17β-carboxylic acid derivative with negligible glucocorticoid activity, than **1** for one from this perspective.^1,2,42^

The rotation of **β** torsion angle affects the position of terminal fluorine atom (F3). In the complex structure GR/**1**, the F3 atom is found to be involved in a weak electrostatic interaction with Asn564 residue (3.84 Å in 3CLD; 3.23 Å in 7PRV),^18,19^ as well as two hydrophobic interactions with Phe749 and Thr739 residues (Figure S3, Supporting Information).^18,19^ The C‒F bond tends to be conformationally flexible, since a weaker density was experimentally detected in this region.^18^ Calculation of the potential energy plots for **β** in **1** and **2**, showed a small rotational barrier (∼2.5 kcal/mol) ranging from 75° to 285° and a large rotational barrier (∼8.7 kcal/mol) in the remaining ranges (Figures 5B and 5E). The **β** in both **1** and **2** behaves flexibly and falls into that range, for example, in **1, β** is 90° in drug formulation state and 309° in the biologically active state;^18^ in **2**, it is 87° in drug formulation state and ∼300° in biologically active state.^18^

Previous literature showed a better fit of 17α-pocket for furoate ring in GR/**1** than propionate ester in GR/**2**, leading to different association and dissociation rates.^18^ Within the 17α-pocket, the furoate ring primarily hydrophobically interacted by Met560, Leu563, Met639 and Met646 residues (Figure S3, Supporting Information).^18,19^ This geometry resulted from the rotation of **γ** in **1**, *i*.*e*., **γ** is 13° in drug formulation state, ∼0° in solution. while it undergoes ∼190° rotations to 204° upon binding in the protein pocket. There is ∼10 kcal/mol rotation barrier in clockwise or anticlockwise direction, that is compensated by the above hydrophobic interactions (Figure 5C). A very weak hydrogen bonding (3.93 Å) between O6 atom of furoate ring and Gln642 residue may also be involved (Figure S3B, Supporting Information).^19^ However, due to the low resolution reported in current X-ray (2.84 Å; PDB entry: 3CLD)^18^ and CryoEM (2.70 Å; PDB entry: 7PRV) structures,^19^ it cannot be verified whether the oxygen atom (O6) should be refined in its current position or the *meta*-position of furoate ring, or if an alternative conformation with **γ** at 24° (180° flipping) co-exists, since it maintains similar interactions but is with minimum conformational changes (15° differences) and energy changes (less than 1.5 kcal/mol; Figure 5C). In contrast, the ethyl part of 17α-propionate ester in **2** within the 17α-pocket is seen to be flexible because of the low rotational barriers (∼1.4 kcal/mol; Figure 5F). Regardless of the oxygen position mentioned above, the large rotational barrier of furoate ring in **1** indicates that it is relatively conformationally fixed in the 17α-pocket, and is reluctant to dissociate; however, the smaller rotational barrier exists in **2** increases entropy upon dissociation with a possible increased tendency towards a release.

In this study, we utilized MicroED to determine the 3D structures of **1** and **2** directly from their microcrystals, which could not be achieved by other structural characterization techniques. These structures are not simply a complement for, or an improvement to the literature, but reveal the different conformations of **1** and **2** in their drug formulation state, pinpointing to various “starting point” before drug functioning in humans. DFT calculations were employed to model the solvent effects and determine the preferred geometries of **1** and **2** in solution, representing the “transition point” before assuming their biologically active state. Finally, we compared structures of **1** and **2** in drug formulation state and in solution with their reported structures in biologically active state to identify the major conformational changes. It was found the steroid backbones are extremely rigid for **1** and **2** in the entire pathway, while significant changes were observed on 17β- and 17α-substitutions, specifically in the **α, β**, and **γ** torsion angles (Figure 4). The potential energy plots for these torsion angles were calculated to estimate their energy landscapes, offering a quantitative approach to understanding their structure-function relationship. It was observed that the conformational changes of 17β-substitution in **2** requires approximately 13 kcal/mol energies to rotate the carbonyl group from backward to an outward position, which is energetically unfavorable compared to less than 1 kcal/mol energy barriers in **1**, suggesting faster association rate of **1** than **2** (Figure 5A and 5D). More unbound 17β-fluoromethylthioester moiety in **2** than **1** metabolized to 17β-carboxylic acid derivative results in shorter biologically half-life.^1,2,42^ While 17α-substitution cannot be metabolized in human, after binding to the 17α-pocket, the furoate ring in **1** is more conformationally rigid (∼10 kcal/mol) than propionate group in **2** (less than 1.4 kcal/mol), suggesting a decreased tendency for dissociation of **1** than **2** (Figure 5C and 5F). This study underscores the combined use of MicroED and DFT calculations to provide a comprehensive understanding of conformational and energetic changes from drug formulation state to biologically active state, explaining how subtle structural differences lead to significant functional changes in pharmaceutical properties.

## Acknowledgements

This study was supported by the National Institutes of Health P41GM136508. Portions of this research or manuscript completion were developed with funding from the Department of Defense MCDC-2202-002. Effort sponsored by the U.S. Government under Other Transaction number W15QKN-16-9-1002 between the MCDC, and the Government. The US Government is authorized to reproduce and distribute reprints for Governmental purposes, notwithstanding any copyright notation thereon. The views and conclusions contained herein are those of the authors and should not be interpreted as necessarily representing the official policies or endorsements, either expressed or implied, of the U.S. Government. The PAH shall flow down these requirements to its sub awardees, at all tiers. The Gonen laboratory is supported by funds from the Howard Hughes Medical Institute.

## Notes

### Competing Interest Statement

The authors have declared no competing interest.

