## Supplementary information for "MicroED Structures of Fluticasone Furoate and Fluticasone Propionate Provide New Insights to Their Function"

### Methods

#### Materials

Fluticasone furoate **1** was commercially purchased from InvivoChem, and fluticasone propionate **2** was commercially purchased from Thermo Scientific Chemicals. Around 8 mg powders of **1** or **2** were separately dissolved in 2 mL methanol. Solvate was allowed to evaporate slowly in room temperature. Needle-shaped microcrystals formed on the surfaces of glass vials and were gently ground to powders using a spatula.

#### Grid preparation

The continuous carbon-coated copper grids (400-mesh, 3.05 mm O.D., Ted Pella Inc.) were pretreated with 15 mA negative glow-discharge plasma for 30 s using PELCO easiGlow (Ted Pella Inc.). The glow-discharged grids were separately mixed with powdery compounds of **1** and **2** in scintillation vials. After 30 s gentle shaking, microcrystals were absorbed on the surface of grids. The grids were clipped outside using c-clips and autogrid rings (Thermo Fisher).

**MicroED data collection**

The autogrids were loaded in a 200 keV Talos Arctica Cryo-TEM (Thermo Fisher) which is equipped with a CMOS CetaD camera (4096 × 4096 pixels) and EPU-D software (Thermo Fisher).[1] The microcrystals were screened under the imaging mode (SA 3400× and 5300×). Since the thickness of samples seriously affects the resolution of electron diffraction, only thin crystals with a certain brightness contrast were selected. Their eucentric heights were carefully calibrated to maintain the crystals inside the beam area during the continuous rotation. The MicroED data was collected in the diffraction mode at the camera length of 659 mm (the calibrated sample-detector distance) and beam size 11 under the parallel beam condition (45.2% C2 intensity). The 70 μm C2 aperture and 50 μm selected area (SA) aperture were used to reduce the background noise, which resulted an approximately 1.4 μm width beam area (electron dose rate: ~0.01 $e^{-1}/(Å^2 \cdot s)$).[2] Typical data collection used a constant rotation rate of 2°/s over an angular wedge of 120° from -60° to +60°, with 0.5 s exposure time per frame (~1 min per dataset), that result a total dose of ~0.6 $e^{-1}/Å^2$ for each item.

**MicroED data processing**

MicroED data was saved in mrc format and converted to smv format using the mrc2smv software (https://cryoem.ucla.edu/microed).[1] The converted frames were indexed and integrated by XDS.[3,4] **1** was indexed with space group P $2_12_12_1$ and only one dataset reached ~96% completeness. **2** was indexed with space group P $2_1$. The common completeness for one single dataset ranged from 61~83%, therefore four datasets were merged to reach ~95% completeness. The merged data was scaled using XSCALE,[4] and their intensities were converted to SHELX hkl format using XDSCONV.[4] Structures of **1** and **2** were *ab initio* solved by SHELXD[5] and refined by SHELXL[6] using Shelxle[7] as a graphical interference (Table S2).

**Density functional theory (DFT) calculation: geometric optimization**

DFT calculations were conducted in ORCA 5.0[8] software using functional/basis set combination B3LYP/6-31G(d,p)[9,10] for geometric optimization. The effects of solvent water were treated using the conductor-like polarizable continuum model (CPCM)[11] and the solvation model based on density (SMD)[12] implemented in ORCA 5.0. The input coordinates were extracted from MicroED structures of **1** and **2** (the minor conformation was omitted). The molecules were allowed to be freely optimized during the calculations. Geometric optimized structures were further compared with other functional/basis set combinations like ωB97X/6-311G(d,p),[13,14] B3LYP/311G(d,p)[9,14] and showed no discernible differences.

**DFT calculation: potential energy calculation**

Three sets of DFT calculation were performed for structure **1** and **2** to evaluate the potential energy changes per bond rotation (see α, β, γ in Figure S2). Take set1 of **1** as an example, α (C13–C17–C20–S1) was rotated from 0° to 360° with 15° increment to generate 25 structures. The coordinates were extracted to generate 25 different input files. α was then fixed while the remaining structure was allowed to freely optimize during the calculations. DFT calculations were conducted within ORCA 5.0,[8] using functional/basis set combination B3LYP/6-31G(d,p),[9,10] and CPCM[11] and SMD[12] models for solvent water effects. The potential energy (relative energy) values were calculated by using the single point energy to substrate the minimum energy. The potential energy versus torsion angle was plotted using GraphPad Prism 8 software (Figure 5).[15]

**Table S1** Summary of unit cell parameters for literature-reported polymorphs (solvomorphs) of **1** and **2**.

| **Solvomorphs of 1** | | | | |
|---|---|---|---|---|
| **1•Guest molecule** | **Space group** | **Unit cell parameters** | **Method** | **Refs** |
| Methy lacetate | $P2_12_12_1$ | a=12.1 Å, b=14.6 Å, c=16.3 Å, β=90° | PXRD | [16] |
| 1,3-dimethylimidazolidinone (DMI) | C2 | a=30.4 Å, b=7.5 Å, c=14.7 Å, β=105.6° | PXRD | [16] |
| (S)-2-Butanol | $P2_12_12_1$ | a=12.4 Å, b=15.5 Å, c=15.5 Å, β=90° | SC-XRD | [16] |
| Ethanol | $P2_12_12_1$ | a=12.2 Å, b=15.2 Å, c=15.5 Å, β=90° | PXRD | [17] |
| Propan-1-ol | $P2_12_12_1$ | a=12.4 Å, b=15.4 Å, c=15.5 Å, β=90° | PXRD | [17] |
| Propan-2-ol | $P2_12_12_1$ | a=12.3 Å, b=15.1 Å, c=15.7 Å, β=90° | PXRD | [17] |
| 1,4-Dioxane | $P2_12_12_1$ | a=12.5 Å, b=14.6 Å, c=16.1 Å, β=90° | PXRD | [17] |
| Ethyl formate | $P2_12_12_1$ | a=12.0 Å, b=14.7 Å, c=16.2 Å, β=90° | PXRD | [17] |
| Acetic Acid | $P2_12_12_1$ | a=11.9 Å, b=14.5 Å, c=16.1 Å, β=90° | PXRD | [17] |
| Acetone | $P2_12_12_1$ | a=11.9 Å, b=14.7 Å, c=16.2 Å, β=90° | PXRD | [17] |
| Dimethylformamide | $P2_12_12_1$ | a=12.1 Å, b=14.8 Å, c=16.2 Å, β=90° | SC-XRD | [17] |
| Dimethylacetamide | $P2_12_12_1$ | a=12.2 Å, b=14.9 Å, c=16.6 Å, β=90° | PXRD | [17] |
| Methylethylketone | $P2_12_12_1$ | a=12.0 Å, b=14.9 Å, c=16.3 Å, β=90° | PXRD | [17] |
| Tetrahydrofuran | $P2_12_12_1$ | a=12.0 Å, b=14.6 Å, c=16.4 Å, β=90° | SC-XRD | [17] |
| N-Methyl-2-pyrrolidinone | $P2_12_12_1$ | a=12.0 Å, b=14.9 Å, c=16.8 Å, β=90° | PXRD | [17] |
| Butan-1-ol | $P2_12_12_1$ | a=12.5 Å, b=15.7 Å, c=15.4 Å, β=90° | PXRD | [17] |
| Methyl acetate | $P2_12_12_1$ | a=12.1 Å, b=14.6 Å, c=16.3 Å, β=90° | PXRD | [17] |
| Toluene | $P2_12_12_1$ | a=7.8 Å, b=13.7 Å, c=34.2 Å, β=90° | PXRD | [18] |
| m-xylene | $P2_12_12_1$ | a=7.8 Å, b=13.8 Å, c=35.9 Å, β=90° | PXRD | [18] |
| Fluorobenzene | $P2_12_12_1$ | a=7.7 Å, b=13.9 Å, c=38.7 Å, β=90° | PXRD | [18] |
| Ethylbenzene | $P2_12_12_1$ | a=7.7 Å, b=13.8 Å, c=37.8 Å, β=90° | PXRD | [18] |
| Chlorobenzene | $P2_12_12_1$ | a=7.8 Å, b=13.7 Å, c=39.3 Å, β=90° | PXRD | [18] |
| Triethylamine | $P2_12_12_1$ | a=7.5 Å, b=12.8 Å, c=34.4 Å, β=90° | SC-XRD | [19] |
| Diethylamine | $P2_12_12_1$ | a=7.7 Å, b=12.7 Å, c=31.9 Å, β=90° | SC-XRD | [19] |
| Dipropylamine | $P2_12_12_1$ | a=7.8 Å, b=13 Å, c=32.2 Å, β=90° | PXRD | [19] |
| **Polymorphs of 2** | | | | |
| **2** | **Space group** | **Unit cell parameters** | **Method** | **Refs** |
| form 1 | $P2_1$ | a=7.7 Å, b=14.2 Å, c=11.3 Å, β=98.5° | PXRD | [20] |
| | $P2_1$ | a=7.6 Å, b=14.1 Å, c=11.0 Å, β=99.3° | SC-XRD | [21] |
| form 2 | $P2_12_12_1$ | a=23.4 Å, b=14.0 Å, c=7.7 Å, β=90° | PXRD | [20] |
| | $P2_12_12_1$ | a=23.2 Å, b=14.0 Å, c=7.7 Å, β=90° | PXRD | [22] |

**Notes**: polymorphs without index of unit cell parameters were omitted for clarification.

**Table S2** MicroED data statistics of **1** and **2**.

| Compound | **1** | **2** |
|---|---|---|
| Name | fluticasone furoate | fluticasone propionate |
| Stoichiometric formula | $C_{27}H_{29}F_3O_6S$ | $C_{25}H_{31}F_3O_5S$ |
| Mr | 538.56 | 500.56 |
| Temperature (K) | 80 | 80 |
| Crystal system | orthorhombic | monoclinic |
| Space group | $P\ 2_1\ 2_1\ 2_1$ | $P\ 2_1$ |
| Unit cell lengths (Å) | | |
| a | 7.70 | 7.57 |
| b | 13.95 | 14.06 |
| c | 23.48 | 10.86 |
| Unit cell angles (°) | | |
| $\alpha$ | 90.0 | 90.0 |
| $\beta$ | 90.0 | 99.4 |
| $\gamma$ | 90.0 | 90.0 |
| Cell volume (Å$^3$) | 2522.10 | 1140.39 |
| No. of datasets merged | 1 | 4 |
| No. of observed reflections | 7217 | 11064 |
| No. of unique reflections | 2051 | 1352 |
| $R_{obs}$ (%) | 16.7 | 23.3 |
| $R_{meas}$ (%) | 19.6 | 24.9 |
| I/Sigma | 6.53 | 7.24 |
| $CC_{1/2}$ | 97.2 | 97.0 |
| Completeness (%) | 96.4 | 94.9 |
| **Resolution (Å)** | **0.90** | **0.96** |
| **$R_1$ (%)** | **16.4** | **15.3** |
| $wR_2$ (%) | 37.1 | 37.9 |
| GooF | 1.290 | 1.341 |

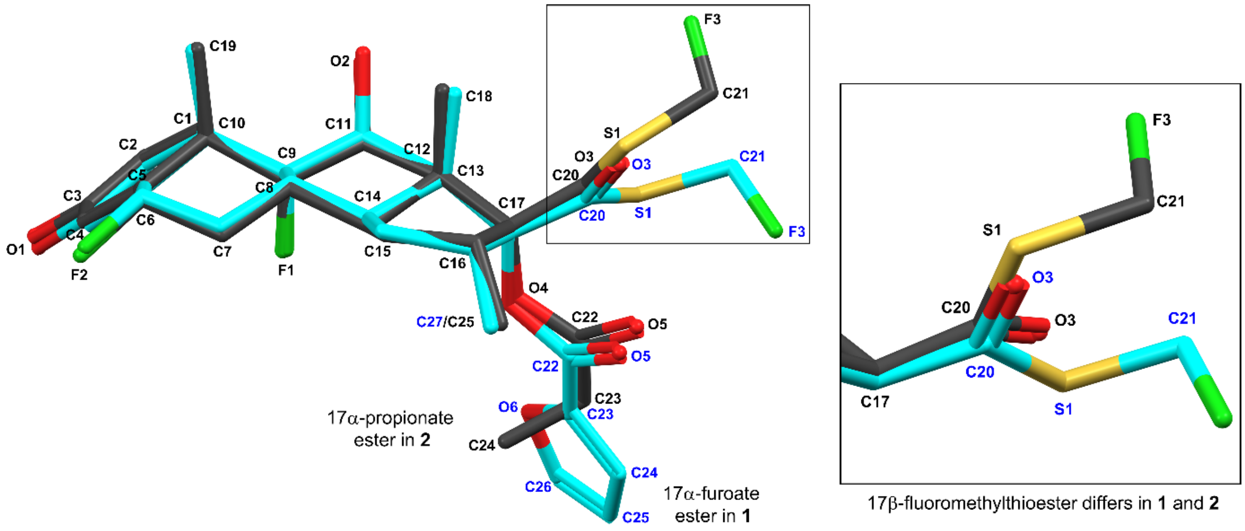


**Figure S1** Overlay of the MicroED structures of **1** and **2**. Carbon atoms of **1** were colored in cyan, carbon atoms of **2** were colored in grey. Numbering of **1** was colored in blue, numbering of **2** was colored in black. Atoms with close locations shared same labels and were colored in black for clarification. 17β-substitutions shown the major conformational changes and were expanded. The minor conformation of **2** was omitted for clarification.

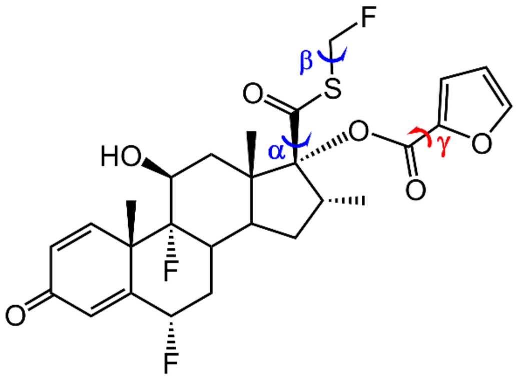


**Fluticasone furoate 1**

α: C13–C17–C20–S1 (set 1)
β: C20–S1–C21–F3 (set 2)
γ: O4–C22–C23–O6 (set 3)

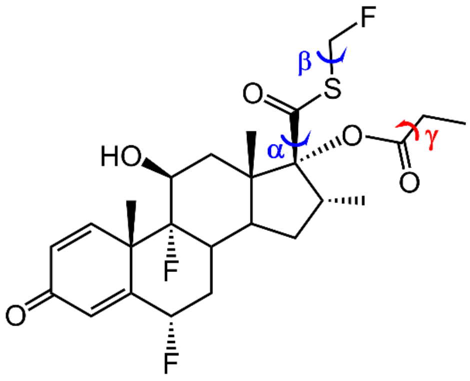


**Fluticasone propionate 2**

α: C13–C17–C20–S1 (set 1)
β: C20–S1–C21–F3 (set 2)
γ: O4–C22–C23–C24 (set 3)

**Figure S2** Structure models and selected torsion angles used for DFT calculations. In sets 1-3 for **1**, **α** (C13–C17–C20–S1), **β** (C20–S1–C21–F3), or **γ** (O4–C22–C23–O6) was rotated and fixed from 0° to 360° with 15° increment, separately. The remaining structure was allowed to freely optimize. In sets 1-3 for **2** (the minor conformation was omitted), **α** (C13–C17–C20–S1), **β** (C20–S1–C21–F3), or **γ** (O4–C22–C23–C24) was rotated and fixed from 0° to 360° with 15° increment, separately. The remaining structure was allowed to freely optimize.

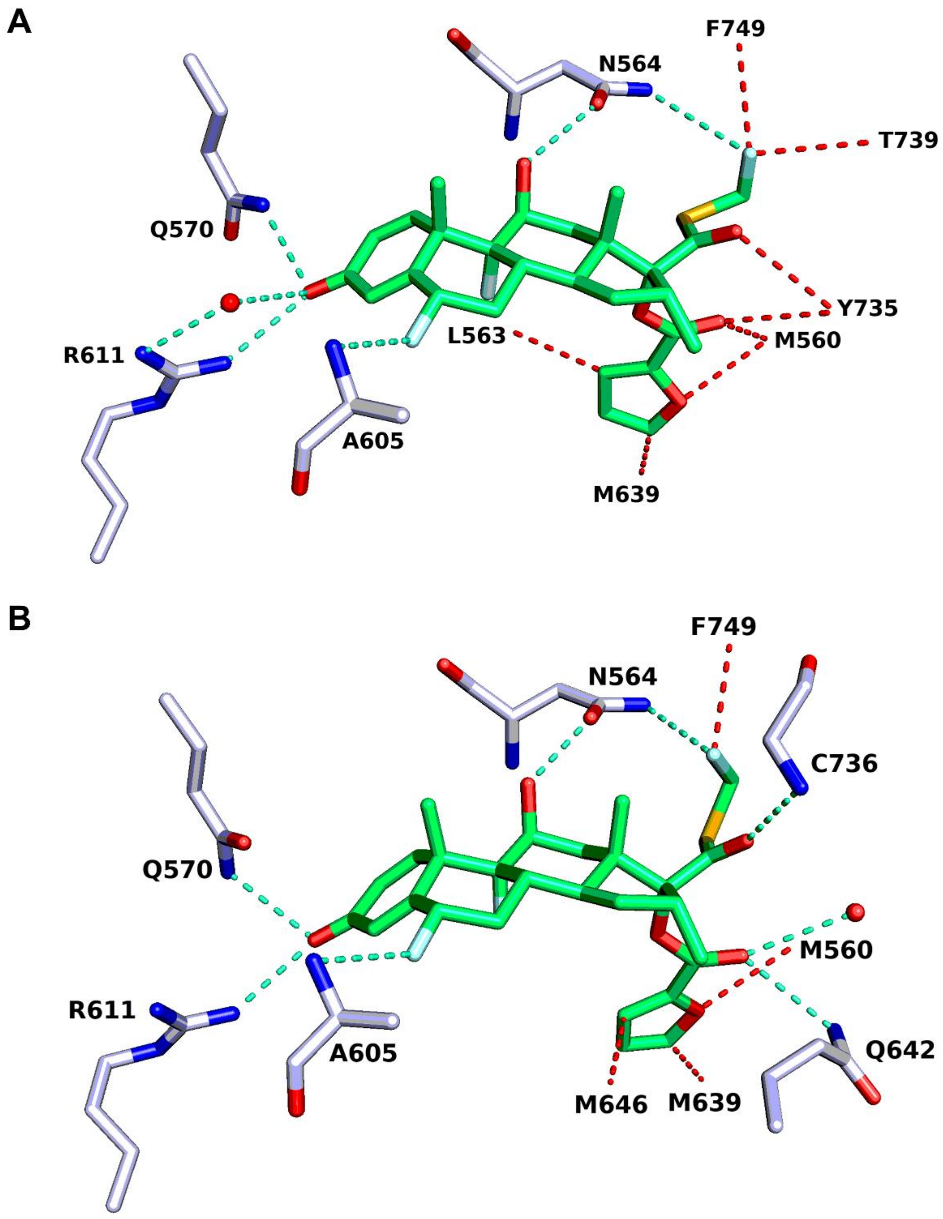


**Figure S3** Interaction between **1** and residues in glucocorticoid receptor (GR). (A) PDB entry: 3CLD;[23] (B) PDB entry: 7PRV.[24] Hydrogen bonding were colored in dashed cyan lines, selected hydrophobic interactions were colored in dashed red lines.
